# Targeting the latent human cytomegalovirus reservoir with virus specific nanobodies

**DOI:** 10.1101/2020.05.12.071860

**Authors:** Timo W.M. De Groof, Elizabeth G. Elder, Raimond Heukers, Eleanor Y. Lim, Mark Wills, John H. Sinclair, Martine J. Smit

## Abstract

Latent reservoirs of viral pathogens are significant barriers to eradication of these viruses. During latency, herpesviruses maintain their genome, with little gene expression, making latent infections refractory to current treatments targeting viral replication. In the case of human cytomegalovirus (HCMV), sporadic reactivation events are well controlled by the immune system. However, in immunocompromised or immunosuppressed individuals, HCMV reactivation often results in morbidity in solid organ and stem cell transplant patients. Clearance of the latent reservoir could lower the incidence and severity of HCMV-associated disease. Here, we develop a virus specific nanobody (VUN100b) that partially inhibits signaling of the viral receptor US28. VUN100b treatment partially reverses latency without fully reactivating the virus. Moreover, VUN100b treatment drives recognition and killing of latently infected monocytes by autologous cytotoxic T lymphocytes from HCMV-seropositive individuals. This study shows the potential of VUN100b as a therapy to clear the HCMV latent reservoir of transplant patients.

## Introduction

Latent reservoirs of viral pathogens are significant barriers to eradication of these viruses from their hosts ^1, 2, 3^. During latency, human herpesviruses and retroviruses maintain their viral genomes in the absence of infectious virus particle production, often with little-to-no viral gene expression ^4^. As such, latent infections are refractory to treatment with typical antivirals that target replication of the virus ^2^. Furthermore, the low number of latently infected cells and the relatively low levels of viral gene expression during latency reduces the levels of viral antigens that would otherwise be readily detectable by the host immune system ^3^. Reactivation from latency results in dissemination and reseeding of the virus. In the case of the ubiquitous betaherpesvirus human cytomegalovirus (HCMV), such sporadic reactivation events are well controlled by a combination of cellular and humoral immunity ^3^. In particular, cytotoxic T lymphocytes (CTLs) against the HCMV immediate early (IE) antigens are present at high frequency in HCMV seropositive individuals ^5^. In immunocompromised or immunosuppressed individuals this control of reactivation is lost, and for both solid organ and stem cell transplant patients, HCMV reactivation frequently results in a disseminated viral infection that is a leading cause of transplant rejection and mortality ^3^.

We have previously proposed that targeting the latent viral reservoir in graft donors and recipients could lower the incidence and severity of HCMV-associated disease in transplant patients ^3^. Latently infected CD34^+^ hematopoietic progenitor cells and their derived CD14^+^ monocytes suppress IE gene expression via the repression of chromatin structure at the viral major immediate early promoter (MIEP) ^6^. By using pan-specific histone deacetylase inhibitors, we previously showed that transient activation of lytic IE gene expression in latently infected monocytes results in CTL-mediated killing of infected cells ^7^. However, we wished to develop a virus-specific molecule that would limit off-target effects but, similarly, induce IE gene expression for use as a shock-and-kill therapeutic.

Here, we show the generation of a bivalent nanobody, VUN100b, from a previously described monovalent nanobody ^8^. VUN100b binds and partially inhibits signaling of the viral protein US28, a chemokine receptor with high homology to human chemokine receptors that is expressed during HCMV latency ^9, 10, 11^. Importantly, a number of reports using particular models of HCMV latency have shown that US28 signaling is essential for the establishment and maintenance of HCMV latency, which is due, at least in part, to US28-mediated repression of the MIEP ^9, 12, 13, 14^. We now show that VUN100b is a partial inverse agonist of US28, resulting in transient activation of the MIEP and subsequent IE gene expression during latency, but no full virus reactivation. We also show that VUN100b drives recognition and killing of latently infected monocytes by CTLs, and demonstrate the efficacy of VUN100b for lowering latent viral loads in *ex vivo* experimentally infected peripheral blood mononuclear cells.

## Results

### Generation of a partial inverse agonistic US28-targeting nanobody

The virally encoded chemokine receptor US28 is absolutely essential for HCMV latency in a number of myeloid cell models of HCMV latency ^9, 12, 13^. Consistent with this, inhibition of US28 using the small-molecule US28 inhibitor VUF2274 resulted in untimely reactivation of the full viral lytic transcription program and production of new infectious viral particles ^13^. However, VUF2274 also showed substantial toxicity and is known to inhibit general CCR1 signaling ^13, 15^. Consequently, we reasoned that new, highly specific reagents that target and inhibit US28 would be needed for any safe shock-and-kill strategy.

To this end, we have developed a new partial inverse agonistic US28-targeting nanobody in the hope that it would inhibit US28 function and efficiently induce IE gene expression for subsequent targeting by host IE-specific CTLs. Nanobodies targeting the extracellular domains of several chemokine receptors are antagonistic as monovalent formats and display inverse agonistic properties as bivalent nanobodies ^16, 17^. We therefore developed a bivalent format of our existing nanobody (VUN100) ^8^ with high affinity for the extracellular domains of US28, which we termed VUN100b. VUN100b was created by fusing two VUN100 molecules using a 30GS linker and showed an approximate 10-fold increase in binding affinity compared to the monovalent VUN100 (0.2 nM ± 0.1 vs 2 nM ± 1) (Figure 1A). Similarly, VUN100b displaced ^125^I-labeled CX3CL1, a known US28 ligand, with approximately 10-fold higher pKi compared to monovalent VUN100 (9.4 ± 0.4 vs 8.1 ± 0.1) (Figure 1B). Next, we tested the functional effect of VUN100b on the constitutive activity of US28 by assessing US28-mediated NFAT (Nuclear Factor of Activated T-cells) activation (Figure 1C). VUN100b inhibited US28 constitutive activity by up to 50% (P value = 0.0005), while no effect was seen for VUN100 or an irrelevant nanobody. By competing for the endogenous ligands of US28 and inhibiting the constitutive activity of US28, VUN100b acts as a partial inverse agonist.

**Fig. 1.**
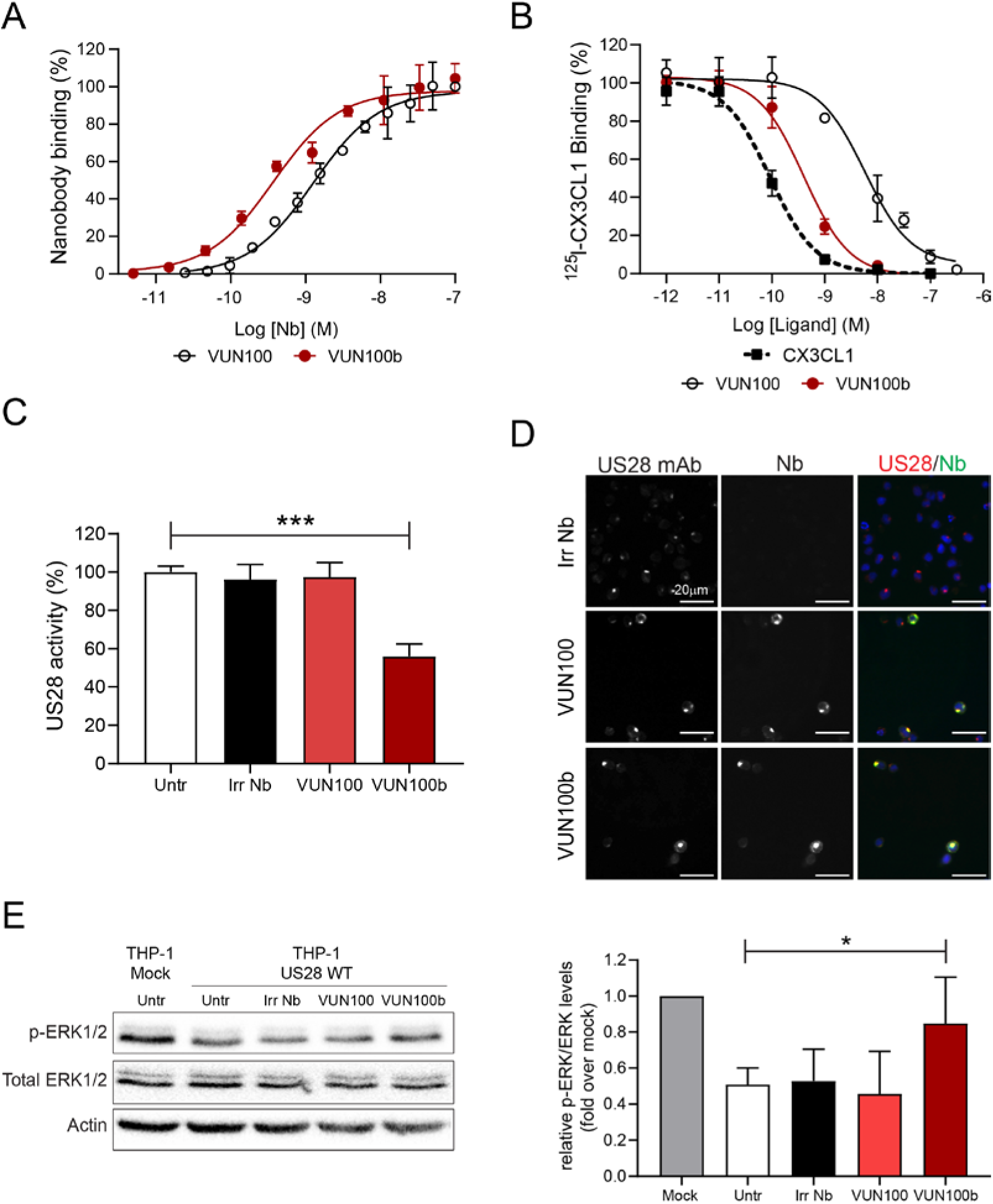
VUN100b binds and inhibits US28 signaling. **A)** ELISA binding of monovalent VUN100 and bivalent VUN100b to membrane extracts of US28-expressing HEK293T cells. **B)** Displacement of ^125^I-CX3CL1 from US28-expressing membranes by unlabeled ligand or the nanobodies VUN100 and VUN100b. **C)** Effect of nanobodies on US28-mediated NFAT (Nuclear Factor of Activated T-cells) activation. HEK293T cells expressing US28 and containing an NFAT-luciferase reporter were untreated (untr) or treated with an irrelevant nanobody (Irr Nb), VUN100 or VUN100b for 24 h prior to luminescence measurement. **D)** Immunofluorescence microscopy of nanobody binding to US28-expressing THP-1 cells. US28 was detected using a polyclonal rabbit anti-US28 antibody (US28 mAb). Bound nanobody was detected using the Myc-tag present on the nanobodies and an anti-Myc antibody (Nb). **E)** Western blot detection for phospho-ERK1/2 of lysates of THP-1 mock transduced cells or US28-expressing THP-1 cells. Cells were untreated (untr) or treated with an irrelevant nanobody (Irr Nb), VUN100 or VUN100b for 48 h. Phosphorylated protein levels over total protein levels were determined and normalized to actin protein levels. Relative phosphorylated protein levels were normalized to untreated THP-1 mock cell lysates. Data is plotted as mean ± S.D.. Statistical analyses were performed using unpaired two-tailed t-test. *, p < 0.05; ***, p < 0.001.

Next, we evaluated the binding to US28 and inverse agonistic activity of VUN100b in the monocytic THP-1 cell line, an established model for HCMV latency ^18^. Both VUN100 and VUN100b bound to US28-expressing THP-1 cells, unlike the irrelevant nanobody (Figure 1D). In addition, none of the three nanobodies bound to mock transduced THP-1 cells, which do not express US28, indicating that these nanobodies are specific to our target (Supplementary Figure 1). We then assessed the effect of the anti-US28 nanobodies on phosphorylation of ERK1/2 (Figure 1E) as US28 is known to attenuate the MAP kinase pathway (ERK1/2) in undifferentiated monocytes in order to suppress MIEP activity during latency ^13^. Treatment of US28-expressing THP-1 cells with VUN100b partially restored the phosphorylation of ERK1/2 (P value = 0.047), indicating a partial inverse agonistic activity of VUN100b in a monocytic cell context. This was not observed for the monovalent VUN100 or irrelevant nanobody. Altogether, our results show that, while both VUN100 and VUN100b can bind to US28, only VUN100b is able to inhibit constitutive US28 signaling in both HEK293T cells and monocytic THP-1 cells.

### US28 nanobodies partially reactivate latently infected CD14^+^ monocytes

Because repression of HCMV MIEP is a downstream consequence of US28 signaling in latently infected myeloid cells, we hypothesized that US28 inhibition by the inverse agonist VUN100b might drive reactivation of viral IE expression from the MIEP in otherwise latently infected cells. Consequently, we determined the effect of the US28 nanobodies on latently infected monocytes. Primary CD14^+^ monocytes were isolated, latently infected with HCMV and treated with nanobodies. At two and six days post infection, IE-expression was assessed (Figure 2A-B and Supplementary Figure 2). As expected, treatment with VUN100b resulted in an increase in IE-expressing monocytes compared to untreated or irrelevant nanobody treated monocytes (Figure 2A-B, P value = 0.0092). Interestingly, we saw a small but significant increase in IE-expression with the antagonistic monovalent VUN100 in 3 out of 4 donors (Figure 2A-B and Figure S2, P value = 0.018). To quantify full reactivation and subsequent virus production, latently infected cells were co-cultured with indicator fibroblasts, a cell type permissive for lytic infection. Importantly, none of the nanobodies, including VUN100b, resulted in any significant plaque formation (Figure 2C). In contrast, production of infectious virus from latently infected monocytes was induced with the phorbol ester PMA (phorbol myristate acetate), which is known to induce myeloid differentiation and reactivation of lytic infection (P value = 0.0095). Taken together, these results indicate that VUN100b treatment results in only a partial reactivation of the lytic transcription programme in latently infected CD14^+^ monocytes. While reactivating the monocytes at the level of IE expression, the nanobody treatments do not result in reactivation of full virus production.

**Fig. 2.**
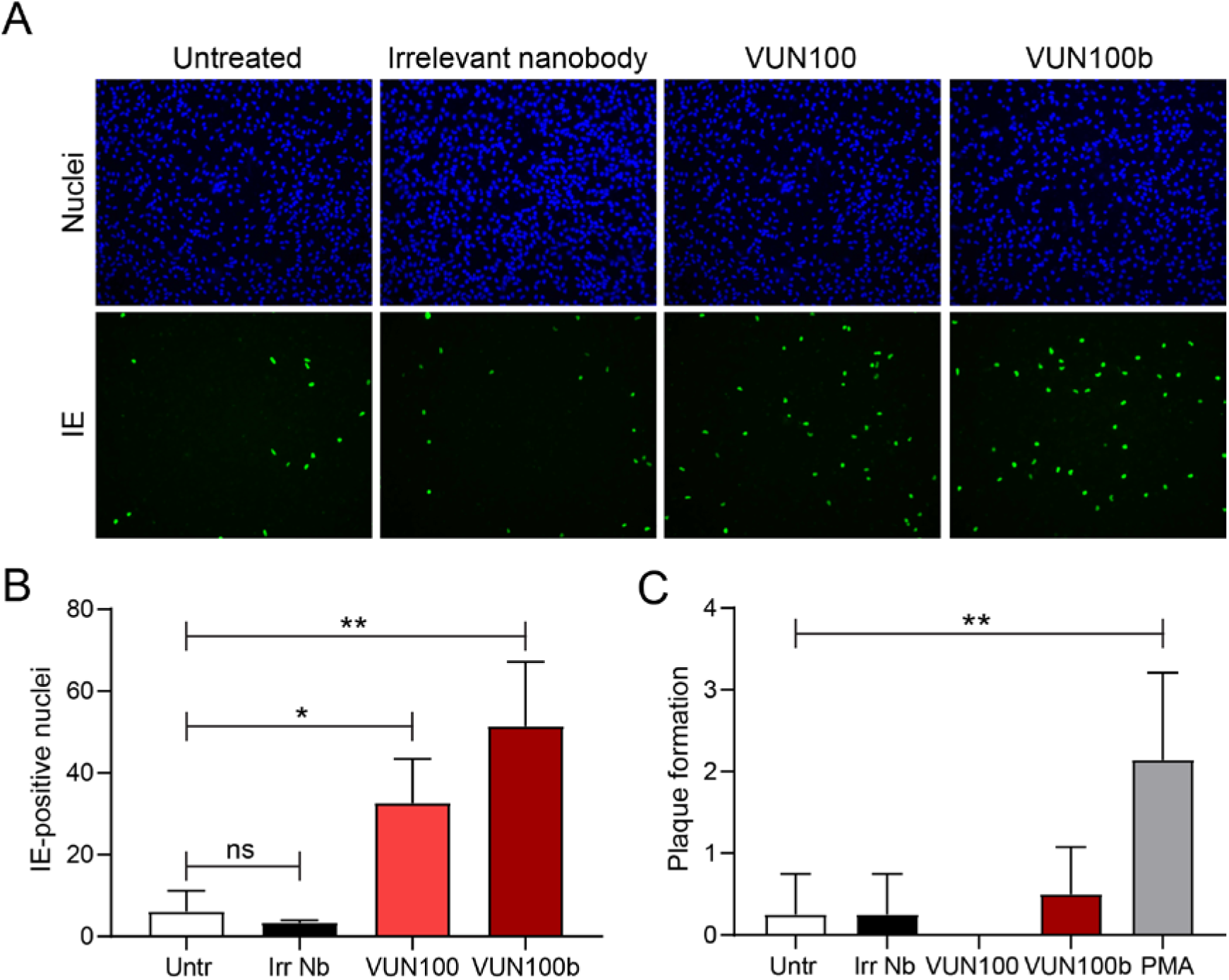
VUN100b induces immediate early expression but no full viral reactivation. **A)** CD14^+^ monocytes were isolated, infected with HCMV IE2-YFP and treated with an irrelevant nanobody or VUN100b. Two days post infection, cells were fixed and stained for immediate early (IE)- expression. **B)** CD14^+^ monocytes were isolated, infected with HCMV IE2-YFP and treated with an irrelevant nanobody (Irr Nb), VUN100 or VUN100b. IE-positive nuclei were counted 6 days post infection. **C)** Six days post infection, nanobody-treated monocytes or monocytes pre-treated with 20 ng/ml PMA before infection (PMA) were co-cultured with Hff1 fibroblasts. Plaque formation was quantified after 4 days of co-culturing. Data is plotted as mean ± S.D.. Statistical analyses were performed using unpaired two-tailed t-test. ns, p > 0.05; *, p < 0.05; **, p < 0.01.

To analyse the extent of reactivated lytic gene expression induced by VUN100b in more detail, we assessed viral gene expression in latently infected monocytes treated with nanobodies. To do this, we used monocytes treated with PMA, which induces myeloid differentiation and reactivation of full lytic infection. We also tested monocytes infected with the ΔUS28 virus, which is devoid of US28 expression and, therefore, undergoes full lytic infection ^13^. At 6 days post infection, RNA was isolated and gene expression of different markers were tested by RT-qPCR (Figure 3). As expected, PMA treatment of infected monocytes or infection with the ΔUS28 virus both resulted in increased levels of transcripts from the major immediate early IE72 gene, the early UL44 gene, the late UL32 gene and the US11 immune evasion gene (Figure 3).

**Fig. 3.**
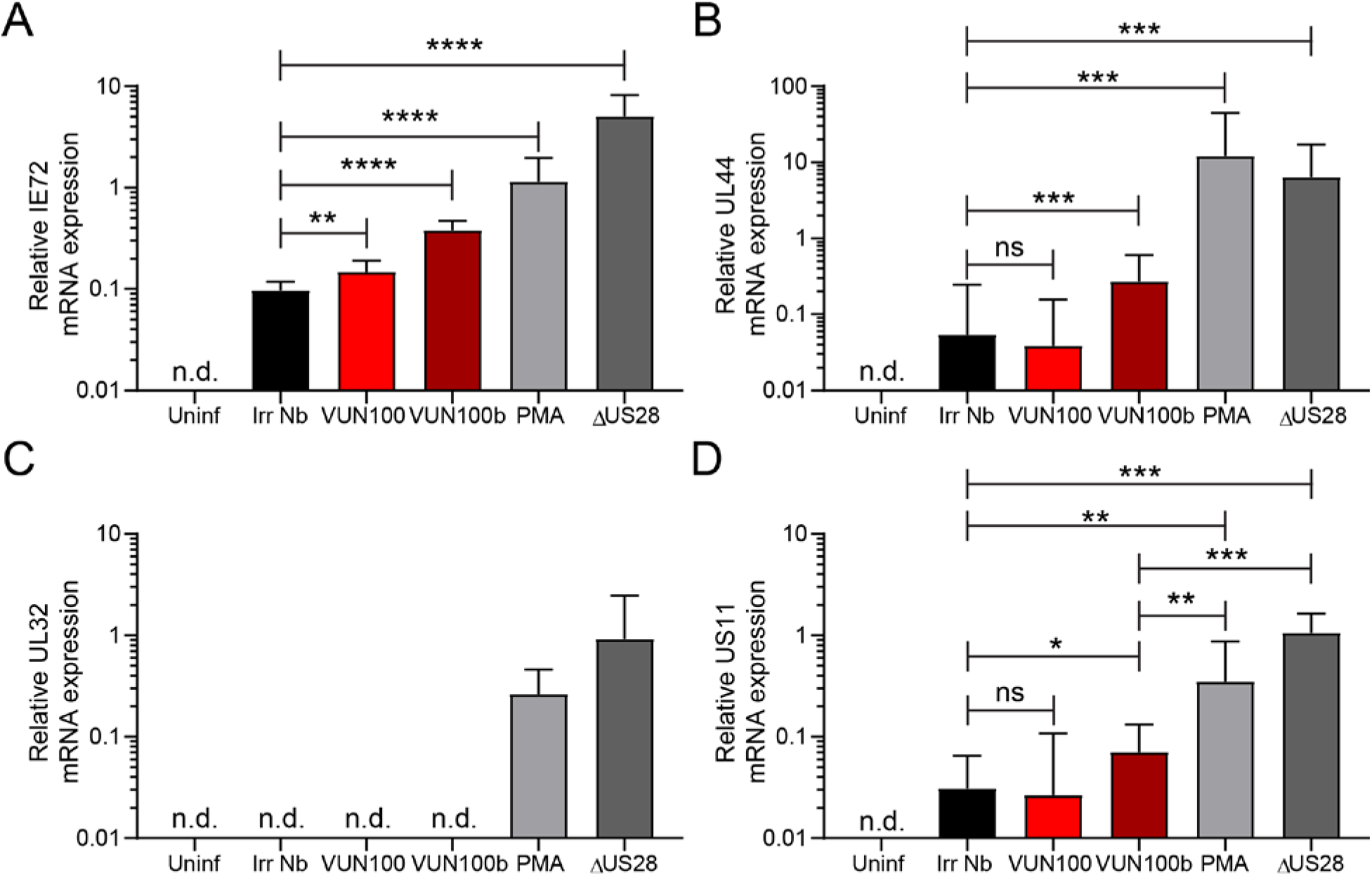
VUN100b increases immediate early, early and late gene but no true late gene expression. CD14^+^ monocytes were uninfected (Uninf), infected with HCMV IE2-YFP or HCMV lacking US28 (ΔUS28). Monocytes were treated with an irrelevant nanobody (Irr Nb), VUN100, VUN100b or pre-treated with 20 ng/ml PMA before infection (PMA). Six days post infection, RNA was isolated and IE72 (**A**), UL44 (**B**), UL32 (**C**) and US11 (**D**) gene expression was measured by RT-qPCR. Data is plotted as mean ± S.D.. Statistical analyses were performed using unpaired two-tailed t-test. n.d.: not detected; ns, p > 0.05; *, p < 0.05; **, p < 0.01; ***, p < 0.001; ****, p < 0.0001.

Consistent with Figure 2, VUN100 (P value = 0.0052) and, in particular VUN100b (P value < 0.0001), treatment of latently infected CD14^+^ monocytes resulted in increased levels of the major immediate early IE72 transcript compared to treatment with the irrelevant nanobody (Figure 3A). In contrast, expression of the early UL44 gene was only slightly increased by VUN100b (P value = 0.0084) compared with irrelevant nanobody or VUN100 treatment, suggesting that viral DNA replication would not be optimal in these cells (Figure 3B). Importantly, VUN100b treatment did not result in UL32 gene expression (Figure 3C), which encodes the true late virion-associated protein pp150. This is consistent with our inability to detect infectious virus production in these cells (see Figure 2).

Finally, we were minded that any shock-and-kill strategy could be thwarted by the expression of viral immune evasins. We therefore also assessed the effect of nanobody treatment on the expression of the immune evasion gene US11. In contrast to PMA and ΔUS28, the slight increase of US11 gene expression (P value = 0.025) was discernibly and significantly lower for the nanobodies compared to fully reactivating cells (P value = 0.0035 and 0.0001, respectively). Taken together, these results confirm that VUN100b treatment results in an increase of IE-expression, only low levels of other viral gene products, and no full virus reactivation.

### Experimentally HCMV-infected CD14^+^ monocytes are targets for HCMV-specific T-cells upon VUN100b treatment

Since HCMV-positive donors have a high frequency of IE-specific cytotoxic T lymphocytes (CTLs) that likely limit virus dissemination from sporadic reactivation events ^5^, we evaluated whether the VUN100b-induced partial reactivation of infected CD14^+^ monocytes would allow clearance of these cells by HCMV-specific CTLs. For this, CD14^+^ monocytes, CD4/CD8^+^ T cells and T cell-depleted peripheral blood mononuclear cells (PBMCs) were isolated from HCMV-positive donors. These CD14^+^ monocytes were infected with HCMV IE2-YFP and treated with an irrelevant nanobody, VUN100b, or PMA. Six days post infection and nanobody or PMA treatment, IE-positive cells were counted. As seen previously, VUN100b treatment (Figure 4A, P value = 0.0006) and pre-treatment with PMA (Supplementary Figure S) resulted in an increase of IE2-YFP-expressing cells. After these 6 days, the infected CD14^+^ monocytes were further split into two groups for subsequent co-culturing with either T cells or T cell-depleted PBMCs. Co-culturing of T cells with VUN100b-treated CD14^+^ monocytes for two days resulted in a significant decrease (P value = 0.0012) of IE-positive CD14^+^ monocytes compared to co-culturing with T cell depleted PBMCs (Figure 4B). The irrelevant nanobody did not affect the number of IE expressing cells when co-cultured with either T cells or the T cell depleted PBMCs, which is consistent with its inability to modulate IE expression.

**Fig. 4.**
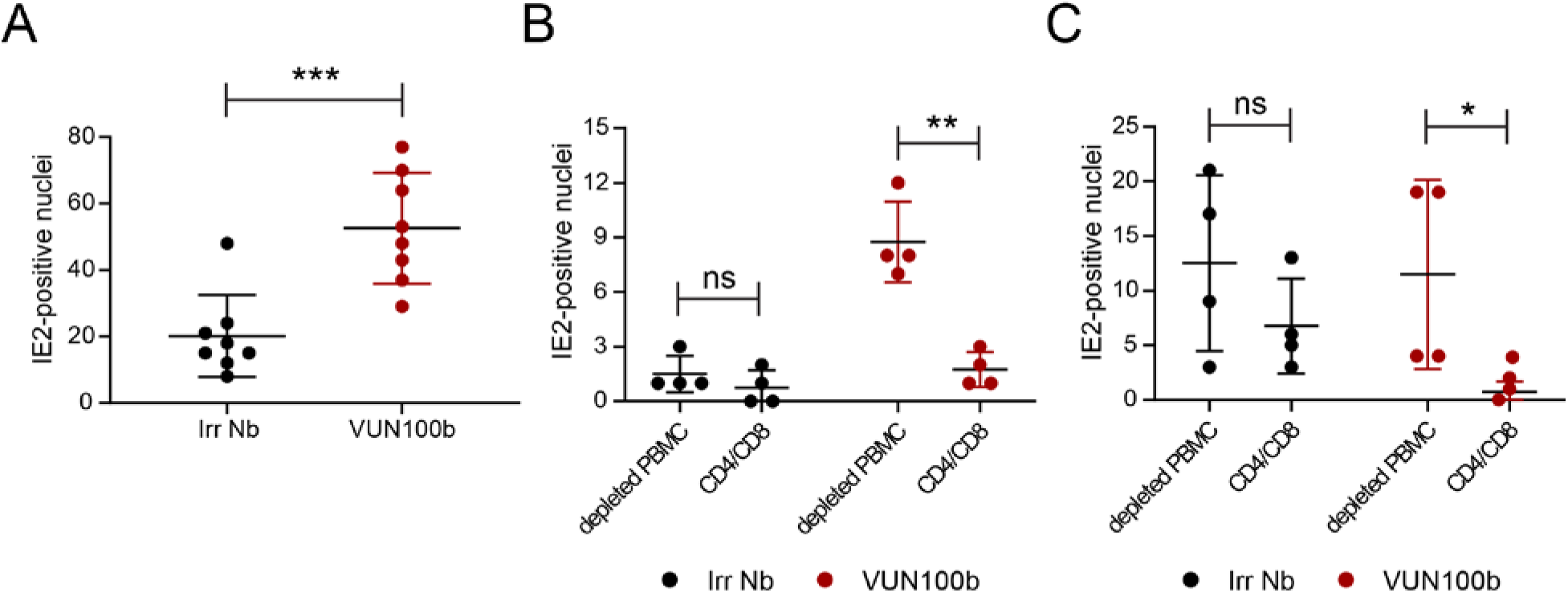
HCMV-infected CD14^+^ monocytes are targets for HCMV specific T-cells upon VUN100b treatment. **A)** Counting of IE2-positive CD14^+^ monocytes before co-culture with CD4/CD8^+^ T cells or T-cell depleted PBMCs. CD14^+^ monocytes were treated with an irrelevant nanobody (Irr Nb) or VUN100b for 6 days post infection. **B)** Counting of IE2-positive CD14^+^ monocytes after 2 days of co-culturing of CD14^+^ monocytes and T cell depleted PBMCs or T cells. **C)** Counting of IE2-positive CD14^+^ monocytes after co-culture of T-cell depleted PBMCs or T cells and differentiation of CD14^+^ monocytes to mature dendritic cells. Data is plotted as mean ± S.D.. Statistical analyses were performed using unpaired two-tailed t-test. ns, p > 0.05; *, p < 0.05; **, p < 0.01; ***, p < 0.001.

After 48 hours of co-culturing, T cells were removed and full reactivation of any remaining HCMV-infected CD14^+^ monocytes was stimulated by differentiating all CD14^+^ monocytes into mature dendritic cells (mDCs) (Figure 4C and Supplementary Figure S). Interestingly, almost no IE-positive mDCs were observed upon treatment with either VUN100b or PMA in combination with T cell co-culturing. This was not seen after co-culturing with T cell depleted PBMCs, indicating a pivotal role for T cells for removal of reactivated cells (P value = 0.049). In contrast, treatment with the irrelevant nanobody did not result in a significant T cell-mediated decrease in IE-positive mDCs.

Although these data nicely illustrate the potential of the VUN100b nanobody, one drawback of the HCMV IE2-YFP virus used in the analysis is that it contains a deletion of the virus US2-US6 region. This region encodes several proteins that interfere with antigen presentation by, for example, downregulating MHC Class I and II molecules ^19, 20, 21^. Though it should be pointed out, that this HCMV IE2-YFP virus does encode US11 (as shown by RT-qPCR, Figure 3D), which can downregulate some MHC Class I molecules ^22, 23^. However, to ensure that our observations would also be recapitulated with a virus with a full complement of immune evasins, we repeated the co-culture experiments described above with a HCMV strain containing an intact US2-6 region. For visualisation and quantification of infection, and in contrast to the IE2-YFP, this virus expresses a C-terminally GFP tagged UL32 protein (Supplementary Figure 4) ^24^. CD14^+^ monocytes from HCMV-positive donors were infected with HCMV UL32-GFP-and treated with VUN100b or an irrelevant nanobody. Six days post infection and nanobody treatment, GFP-positive cells were counted (Supplementary Figure 4A). In line with our observations showing that VUN100b induces IE gene expression but does not induce late lytic gene expression (Figure 3C), no significance difference in the number of UL32-GFP positive cells was seen between the cells treated with the irrelevant nanobody or VUN100b. Upon 48 hours of co-culturing of the infected CD14^+^ monocytes with T cells from HCMV positive donors, T cells were removed, all CD14^+^ monocytes were differentiated into mature dendritic cells. To assess viral load, these dendritic cells were co-cultured with Hff1 fibroblast cells. After 7 days of co-culturing HCMV DNA genome levels were determined (Supplementary Figure 4B). In line with the results obtained with the IE2-YFP virus, significant reduction (P value = 0.0043) of HCMV DNA levels was observed for the VUN100b-treated CD14^+^ monocytes compared to the monocytes treated with the irrelevant nanobody. Overall, these data indicate that partial reactivation of latently infected cells by VUN100b is sufficient to induce IE-expression to levels that allow HCMV-specific T cell-mediated clearing of latently infected monocytes.

## Discussion

HCMV establishes a latent infection in CD34^+^ progenitor cells and CD14^+^ monocytes ^25, 26^. Although only present in a small percentage of these cells, reactivation of HCMV can lead to disease or mortality in immunosuppressed transplant patients and immunocompromised individuals ^27, 28^. Importantly, no current antiviral agents target this latent reservoir, leaving an unmet need to target these latently infected cells. In this study, we set out to target the HCMV-encoded chemokine receptor US28 using a newly developed bivalent US28 nanobody to partially reactivate HCMV-latently infected cells and drive their T-cell recognition and killing.

We used the previously described US28-targeting nanobody VUN100 to develop a new bivalent format VUN100b ^8^. Interestingly, the coupling of two antagonistic monovalent VUN100 nanobodies resulted in a bivalent format with partial inverse agonistic properties. Although the mechanism behind this still has yet to be explained, similar observations were made with other nanobodies targeting chemokine receptors ^29, 30, 31^. VUN100b could bind and partially inhibit US28 signaling in cultured cell lines, and we recapitulated our observations in experimental latency settings in primary CD14^+^ monocytes. Here, we saw upregulation of IE gene expression without substantial immune evasin gene expression, late gene expression or full virus reactivation. This is a major advantage for a shock-and-kill strategy, which requires detection by host immunity ^6, 32^. On a molecular level, our findings suggest that there is a threshold of inhibition of US28 signaling required for full viral reactivation.

Interestingly, the monovalent nanobody VUN100 also induced some IE gene expression in most donors despite not inhibiting US28 constitutive signaling. VUN100 blocks ligand binding, and the ligand binding activity of US28 has been shown to be required for the establishment in some, but not all, experimental latency systems ^8, 12, 13, 33^. Furthermore, the donor-to-donor variability we observed in the VUN100 effect raises questions about the role of US28 ligand binding during HCMV latency in patients. In contrast, VUN100b consistently and robustly induced IE expression in all donors tested and led to recognition and killing by CTLs from seropositive individuals. Because our results were obtained using experimentally latently infected cells, follow up experiments using *ex vivo* PBMCs from naturally infected donors would provide valuable confirmation of the current results. Nevertheless, our current studies are consistent with previous reports of shock-and-kill strategies using epigenetic modifiers, such as histone deacetylase (HDAC) inhibitors, to induce transient viral gene expression ^7, 34, 35, 36^. However, treatment with HDAC inhibitors are associated with substantial off-target effects due to other and more physiological functions of these enzymes ^37^. In contrast to such inhibitors, we have, here, developed a molecule specific to HCMV-infected cells and, therefore, it should display limited off-target effects. Recently, the first FDA-approved nanobody has entered the clinic for treatment of acquired Thrombotic Thrombocytopenic Purpura ^38, 39^. This paves the way for potential therapeutic use of HCMV-specific nanobodies. In addition to their intrinsic activity, the efficacy of such nanobodies in experimental and clinical settings could be further enhanced through coupling to effector molecules ^8, 40, 41, 42, 43, 44^.

Overall, our study provides a strong basis for using inverse agonistic/inhibitory anti-US28 nanobodies to reduce latent viral loads in transplant donors and/or recipients prior to surgery and immunosuppression. This could lead to a lower incidence of CMV-associated disease and mortality during life-saving solid organ and stem cell transplantation.

## Materials and Methods

### Cell culture and virus infection

Primary CD14^+^ monocytes were isolated from apheresis cones (NHS Blood and Transfusion Service, United Kingdom), as described previously^45^, or by magnetic-activated cell sorting (MACS) separation using CD14 microbeads (Miltenyi Biotec, Bergisch Gladbach, Germany) as described previously ^46^. The monocytes were adhered to tissue culture dishes (Corning, Tewksbury, MA, USA). CD4^+^ and CD8^+^ T cells were isolated by MACS from monocyte-depleted PBMC using CD4 and CD8 microbeads (Miltenyi Biotec). These primary cells were cultured in X-vivo 15 (Lonza, Walkersville, MD, USA) supplemented with 2 mM L-glutamine (Gibco, Thermo Fisher Scientific, Waltham, MA, USA) at 37°C in 5% CO_2_. Phorbol myristate acetate (PMA, Sigma-Aldrich, Saint-Louis, MO, USA) was used as described in figure legends at 20 ng/mL.

THP-1 cells, lentivirally transduced with different US28 constructs, have been described previously^13^ and were cultured according to ATCC standards (RPMI-1640 media (Sigma-Aldrich)) supplemented with 10% heat-inactivated fetal bovine serum (FBS; PAN Biotech, Aidenbach, Germany), 100 U/mL penicillin and 100 µg/mL streptomycin (Sigma-Aldrich), and 0.05 mM 2-mercaptoethanol (Gibco) maintained at 37°C in 5% CO_2_).

HEK293T cells were purchased from ATCC (Wesel, Germany) and were grown at 5% CO2 and 37 °C in Dulbecco’s Modified Eagle’s Medium (DMEM, Thermo Fisher Scientific) supplemented with 1% Penicillin/Streptomycin (Thermo Fisher Scientific) and 10% FBS (Thermo Fisher Scientific).

Human foreskin fibroblasts (Hff1; ATCC® SCRC-1041™) were maintained in DMEM (Sigma-Aldrich) supplemented with 10% heat-inactivated FBS and 100 U/mL penicillin and 100 µg/mL streptomycin.

Viral isolate RV1164 (HCMV TB40/E strain with an IE2-YFP tag) has been described previously ^47^. Titan WT and ΔUS28 strains have been described previously ^48^. TB40-UL32-GFP-HCMV has been described earlier ^24^. CD14^+^ monocytes were infected at a multiplicity of infection (MOI) of 3 for 2 hours, were washed twice with phosphate buffered saline (PBS), before replacing with fresh X-vivo 15 + L-glutamine.

### Nanobody production

Nanobody gene fragments were recloned in frame with a myc-His6 tag in the pET28a production vector. Bivalent formats of VUN100 were constructed by addition of a 30GS-linker in frame with the nanobody fragments. Nanobodies were produced as described previously ^49^. Purity of the nanobodies was verified by sodiumdodecyl sulfate-polyacrylamide gel electrophoresis (SDS-PAGE) (Bio-Rad, Hercules, CA, USA).

### Nanobody binding ELISA

Nanobody binding was performed as described previously ^8^. Briefly, US28-expressing membrane extracts were coated in a 96 well MicroWell™ MaxiSorp™ flat bottom plate (Sigma-Aldrich) overnight at 4 °C. Next day, wells were washed and blocked with 2% (w/v) skimmed milk (Sigma-Aldrich) in PBS. Different concentrations of nanobodies were incubated. Nanobodies were detected with mouse-anti-Myc antibody (1:1000, clone 9B11, Cell Signaling Technology, Leiden, The Netherlands) and Goat anti-Mouse IgG-HRP conjugate (1:1000, #1706516, Bio-Rad). Optical density was measured at 490 nm with a PowerWave plate reader (BioTek, Winooski, VT, USA). Data was analyzed using GraphPad Prism version 8.0 (GraphPad Software, Inc., La Jolla, CA, USA).

### Competition binding

Membrane extracts of HEK293T and HEK293T overexpressing US28 were used during competition binding studies. The experiments were performed as described previously ^50^. Data was analyzed using GraphPad Prism version 8.0.

### NFAT reporter gene assay

HEK293T cells were transfected with 50 ng pcDEF3-HA-US28 VHL/E, 2.5 μg NFAT-reporter gene vector (Stratagene, La Jolla, CA, USA) and 2.5 μg empty pcDEF3 DNA as described previously ^8^. Six hours post-transfection, cells were trypsinized using Trypsin-EDTA 0.05% (Gibco) and 50.000 cells were seeded per well in a Poly-L-lysine (Sigma-Aldrich) treated white bottom 96-well assay plate. Nanobodies were added with a final concentration of 100 nM and cells were incubated at 37°C and 5% CO2. After 24h, medium was removed and 25μL LAR (0.83mM D-Luciferine, 0.83mM ATP, 0.78μM Na2HPO4, 18.7mM MgCl2, 38.9mM Tris-HCl (pH 7.8), 2.6μM DTT, 0.03% Triton X-100 and 0.39% Glycerol) was incubated for 30 min at 37°C. Luminescence (1s per well) was measured using a Clariostar plate reader (BMG Labtech, Ortenberg, Germany). Data was analyzed using GraphPad Prism version 8.0.

### Immunofluorescence microscopy

THP-1 cells were spun down at 500x g for 5 min, resuspended in 4% paraformaldehyde (Sigma-Aldrich) and seeded in a 96 well U-bottom plate. Cells were fixed for 10minutes at room temperature. After fixation, cells were permeabilized with 0.5% NP-40 (Sigma-Aldrich) for 30 min at room temperature. Nanobodies were incubated for 1 h at RT and detected using Mouse-anti-Myc antibody (1:1000, 9B11 clone, Cell Signaling). US28 was visualized with the rabbit-anti-US28 antibody (1:1000, Covance, Denver, PA, USA, ^51^). Subsequently, cells were washed and incubated with Goat-anti-Rabbit Alexa Fluor 546 (1:1000 in 1% (v/v) FBS /PBS, Thermo Fisher Scientific) and Goat-anti-Mouse Alexa Fluor 488 (1:1000 in 1% (v/v) FBS/PBS, Thermo Fisher Scientific).

Immediate early antigen was detected in monocytes and fibroblasts by fixation and permeabilization in 70% ethanol at -20 °C for 30 minutes and blocking using PBS with 1 % bovine serum albumin and 5% goat serum, and then incubating with mouse anti-IE antibody (1:1000, #11-003, Argene, bioMériux, Marcy-l’étoile, France,) followed by secondary antibodies as described above.

### Western blot

Mock transduced or US28-expressing THP-1 cells were seeded in a 6 wells plate and incubated with 100 nM nanobodies. After 48 h, cells were lysed in native lysis buffer (25 mM Tris HCL pH7.4, 150 mM NaCl, 1 mM EDTA, 1% NP-40, 5% Glycerol, 1 mM NaF, 1 mM NaVO_3_, cOmplete™ protease inhibitor cocktail) for 10 min on ice. Cell debris was removed by centrifugation at 13,000g. Protein concentration of lysates was determined by Pierce™ BCA protein assay kit (Thermo Fisher Scientific) and same protein quantities were separated on a 10% SDS-PAGE gel under reducing conditions and transferred to 0.45 μm PVDF blotting membrane (GE healthcare, Chicago, IL, USA). Total ERK1/2 and phospho-ERK1/2 were detected using p44/42 MAPK antibody (1:1000 in 5% BSA/TBS-T, #9102, Cell Signaling) and phospho-p44/42 MAPK (Thr202/Tyr204) (1:1000 in 5% BSA/TBS-T, #9106, Cell Signaling). Actin was detected using anti-actin antibody (1:2000 in 5% BSA/TBS-T, Clone AC-74, Sigma-Aldrich). Antibodies were detected using Goat anti-Rabbit IgG-HRP conjugate (1:10000, #1706515, Bio-Rad) or Goat anti-Mouse IgG-HRP conjugate (1:10000, #1706516, Bio-Rad). Blots were developed using Western Lightning Plus-ECL (Perkin-Elmer, Waltham, MA, USA) and visualized with Chemidoc™ (Bio-Rad).

### Detection of immediate early expression

CD14^+^ monocytes were isolated and seeded in a 96 wells plate. As a positive control, CD14^+^ monocytes were pre-treated with 20ng/ml PMA one day after seeding. The next day, medium was removed and cells were infected with RV1164 viral isolate. Two hours post infection, medium was aspirated and replaced with medium containing nanobodies at a final concentration of 100 nM. Three days post infection, nanobody-containing medium was refreshed. IE-expression was detected by means of IE2-YFP tag or staining of IE as described above.

### RNA and DNA extraction and analysis

CD14^+^ monocytes were isolated and infected as described above. 6 days post infection, cells were washed once with 1x PBS and RNA was harvested by adding Trizol reagent (Zymo Research, Irvine, CA, USA). RNA was isolated using Direct-Zol RNA MiniPrep kit (Zymo Research) according to the manufacturer’s instructions. cDNA was produced by Quantitect Reverse Transcription kit (Qiagen, Hilden, Germany) according to manufacturer’s instructions. qPCR was performed using LUNA SYBR green qPCR reagents (New England Biolabs, Ipswich, MA, USA) using primers in the table. Viral transcript levels were normalized to GAPDH and are presented as 2^ΔCt^.

For analysis of viral genomes, cells were washed with PBS, then solution A (100mM KCl, 10mM Tris-HCl pH8.3 2.5mM MgCl) was added followed by an equal volume of solution B (10mM Tris-HCl pH8.3, 2.5mM MgCl 1% Tween20, 1% NP-40, 0.4 mg/ml proteinase K). Cells were scraped and placed in microtubes before heating at 60°C for one hour then 95°C for ten minutes. Two μl of each solution was used in qPCR analysis of the UL44 non-transcribed promoter region, using the GAPDH non-transcribed promotor region to correct for total DNA levels. Primer sequences are provided in the table.

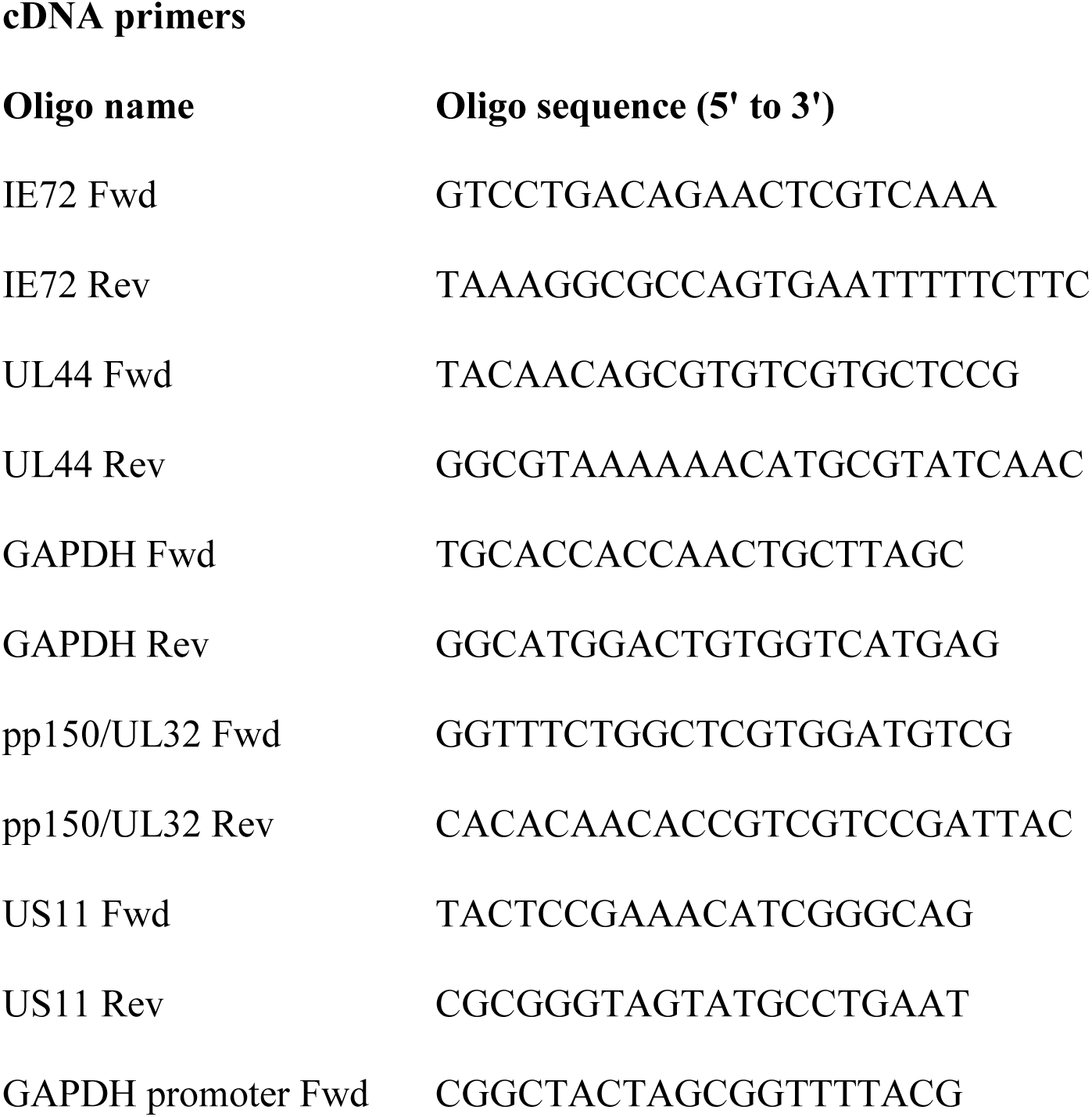
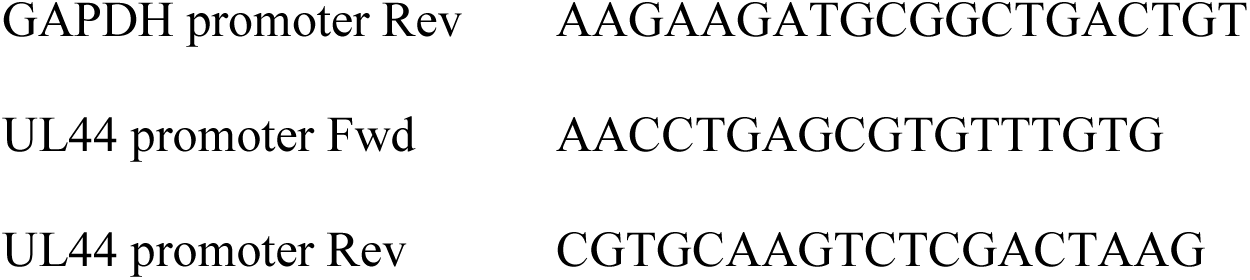

### PBMC and T cell co-culture and virus reactivation

For experimental latency experiments, following PBMC isolation from a HCMV-positive healthy donor peripheral blood, CD14^+^ monocytes were isolated, plated on 96 well plates, and treated as described above and the remained PBMC were frozen in liquid nitrogen until one day prior to co-culture. At this time, the PBMC were thawed and rested overnight. CD4^+^ and CD8^+^ T cell fractions were isolated as above and pooled, and these, or the remaining PBMC were added to the monocyte cultures at an effector:target cell ratio of 5:1. After two days, the T cells/depleted PBMC were washed away using PBS + 2 mM EDTA, and the medium on the monocytes was replenished with X-vivo 15 + L-glutamine containing interleukin-4 (Miltenyi Biotec) and granulocyte-macrophage colony-stimulating factor (Miltenyi Biotec) at 1,000 U/ml in order to stimulate differentiation to immature dendritic cells, along with 10 μg/mL anti HLA-A, B, C (Biolegend) and 10 μg/mL anti HLA-DR, DP, DQ (BD Bioscience) to block any further T cell killing. After 5 days, this medium was aspirated and replaced with X-vivo 15 + L-glutamine supplemented with 50 ng/mL lipopolysaccharide (LPS, Invivogen, San Diego, CA, USA) for 2 days to induce maturation of dendritic cells. This media was aspirated and 5 × 10^3^ Hff1 cells were added to each well in DMEM-10 medium. After 7 days of co-culture, the cells were harvested for viral genome, as described above.

### Ethical approval for the use of human samples

Human samples were obtained under ethical approval from Cambridgeshire 2 Research Ethics Committee (REC reference 97/092) conducted in accordance with the Declaration of Helsinki. All volunteers have given informed written consent before providing blood samples.

### Statistical analysis

For the validation of the nanobody, each experiment is consisting of two to four technical replicates of three biological replicates. For the virus experiments, three to six technical replicates of two to four different donors has been used. No outliers were removed from the experiments. Data were analysed using GraphPad Prism version 8.0 software. All figures are representative figures and all data is plotted as mean ± S.D. Statistical analyses were performed using unpaired two-tailed t-test. P value < 0.05 was considered statistically significant.

### Biological material availability

US28-targeting nanobodies, described in this paper, can be obtained through an MTA. The HCMV strains RV1164 (TB40-UL32-GFP-HCMV), Titan wildtype, Titan ΔUS28 and TB40-UL32-GFP-HCMV are all published and not ours to make available

### Data and materials availability

All data associated with this study are present in the paper or in the Supplementary Materials. A reporting summary for this study is available as a Supplementary Information file. The source data are provided as a Source Data file.

## Supporting information

Supplemental Figures_De Groof, Elder et al_2020

## Acknowledgments

We thank Linda Teague, Roy Whiston and Georgina Brown for technical assistance. This work was supported by the Netherlands Organization for Scientific Research (NWO: Vici grant 016.140.657), the Wellcome Trust (Grant 109075/Z/15/A), the British Medical Research Council (grant MR/K021087/1) and Cambridge NIHR BRC Cell Phenotyping Hub.

## Author contributions

Conceptualization: T.D.G., E.E., R.H., J.S. and M.S.; Investigation: T.D.G, E.E. and E.L.; Formal analysis: T.D.G. and E.E.; Writing-original draft preparation: T.D.G. and E.E; Writing-review and editing: R.H., M.W., J.S and M.S.; Visualization: T.D.G. and E.E; Supervision: R.H., M.W., J.S and M.S.; Funding acquisition: J.S., M.W., and M.S.

## References

1. Sadowski I, Hashemi FB. Strategies to eradicate HIV from infected patients: elimination of latent provirus reservoirs. Cell Mol Life Sci, (2019).

2. Nehme Z, Pasquereau S, Herbein G. Control of viral infections by epigenetic-targeted therapy. Clin Epigenetics 11, 55 (2019).

3. Wills MR, Poole E, Lau B, Krishna B, Sinclair JH. The immunology of human cytomegalovirus latency: could latent infection be cleared by novel immunotherapeutic strategies? Cell Mol Immunol 12, 128–138 (2015).

4. Knipe DM, Raja P, Lee J. Viral gene products actively promote latent infection by epigenetic silencing mechanisms. Curr Opin Virol 23, 68–74 (2017).

5. Khan N, Cobbold M, Keenan R, Moss PA. Comparative analysis of CD8+ T cell responses against human cytomegalovirus proteins pp65 and immediate early 1 shows similarities in precursor frequency, oligoclonality, and phenotype. J Infect Dis 185, 1025–1034 (2002).

6. Elder E, Sinclair J. HCMV latency: what regulates the regulators? Med Microbiol Immunol 208, 431–438 (2019).

7. Krishna BA, Lau B, Jackson SE, Wills MR, Sinclair JH, Poole E. Transient activation of human cytomegalovirus lytic gene expression during latency allows cytotoxic T cell killing of latently infected cells. Sci Rep 6, 24674 (2016).

8. De Groof TWM, et al. Nanobody-Targeted Photodynamic Therapy Selectively Kills Viral GPCR-Expressing Glioblastoma Cells. Mol Pharm 16, 3145–3156 (2019).

9. Humby MS, O’Connor CM. Human Cytomegalovirus US28 Is Important for Latent Infection of Hematopoietic Progenitor Cells. J Virol 90, 2959–2970 (2015).

10. Beisser PS, Laurent L, Virelizier JL, Michelson S. Human cytomegalovirus chemokine receptor gene US28 is transcribed in latently infected THP-1 monocytes. J Virol 75, 5949–5957 (2001).

11. Hargett D, Shenk TE. Experimental human cytomegalovirus latency in CD14+ monocytes. Proc Natl Acad Sci U S A 107, 20039–20044 (2010).

12. Krishna BA, Humby MS, Miller WE, O’Connor CM. Human cytomegalovirus G protein-coupled receptor US28 promotes latency by attenuating c-fos. Proc Natl Acad Sci U S A 116, 1755–1764 (2019).

13. Krishna BA, Poole EL, Jackson SE, Smit MJ, Wills MR, Sinclair JH. Latency-Associated Expression of Human Cytomegalovirus US28 Attenuates Cell Signaling Pathways To Maintain Latent Infection. MBio 8, (2017).

14. Zhu D, et al. Human cytomegalovirus reprogrammes haematopoietic progenitor cells into immunosuppressive monocytes to achieve latency. Nat Microbiol 3, 503–513 (2018).

15. Casarosa P, et al. Identification of the first nonpeptidergic inverse agonist for a constitutively active viral-encoded G protein-coupled receptor. J Biol Chem 278, 5172–5178 (2003).

16. De Groof TWM, Bobkov V, Heukers R, Smit MJ. Nanobodies: New avenues for imaging, stabilizing and modulating GPCRs. Mol Cell Endocrinol 484, 15–24 (2019).

17. Heukers R, De Groof TWM, Smit MJ. Nanobodies detecting and modulating GPCRs outside in and inside out. Curr Opin Cell Biol 57, 115–122 (2019).

18. Lau B, et al. Human cytomegalovirus miR-UL112-1 promotes the down-regulation of viral immediate early-gene expression during latency to prevent T-cell recognition of latently infected cells. J Gen Virol 97, 2387–2398 (2016).

19. Noriega V, Redmann V, Gardner T, Tortorella D. Diverse immune evasion strategies by human cytomegalovirus. Immunol Res 54, 140–151 (2012).

20. Wiertz EJ, Jones TR, Sun L, Bogyo M, Geuze HJ, Ploegh HL. The human cytomegalovirus US11 gene product dislocates MHC class I heavy chains from the endoplasmic reticulum to the cytosol. Cell 84, 769–779 (1996).

21. Johnson DC, Hegde NR. Inhibition of the MHC class II antigen presentation pathway by human cytomegalovirus. Curr Top Microbiol Immunol 269, 101–115 (2002).

22. Zimmermann C, et al. HLA-B locus products resist degradation by the human cytomegalovirus immunoevasin US11. PLoS Pathog 15, e1008040 (2019).

23. Ameres S, Besold K, Plachter B, Moosmann A. CD8 T cell-evasive functions of human cytomegalovirus display pervasive MHC allele specificity, complementarity, and cooperativity. J Immunol 192, 5894–5905 (2014).

24. Sampaio KL, Cavignac Y, Stierhof YD, Sinzger C. Human cytomegalovirus labeled with green fluorescent protein for live analysis of intracellular particle movements. J Virol 79, 2754–2767 (2005).

25. Taylor-Wiedeman J, Sissons JG, Borysiewicz LK, Sinclair JH. Monocytes are a major site of persistence of human cytomegalovirus in peripheral blood mononuclear cells. J Gen Virol 72 (Pt 9), 2059–2064 (1991).

26. Hahn G, Jores R, Mocarski ES. Cytomegalovirus remains latent in a common precursor of dendritic and myeloid cells. Proc Natl Acad Sci U S A 95, 3937–3942 (1998).

27. Slobedman B, Mocarski ES. Quantitative analysis of latent human cytomegalovirus. J Virol 73, 4806–4812 (1999).

28. Sissons JG, Wills MR. How understanding immunology contributes to managing CMV disease in immunosuppressed patients: now and in future. Med Microbiol Immunol 204, 307–316 (2015).

29. Jahnichen S, et al. CXCR4 nanobodies (VHH-based single variable domains) potently inhibit chemotaxis and HIV-1 replication and mobilize stem cells. Proc Natl Acad Sci U S A 107, 20565–20570 (2010).

30. Maussang D, et al. Llama-derived single variable domains (nanobodies) directed against chemokine receptor CXCR7 reduce head and neck cancer cell growth in vivo. J Biol Chem 288, 29562–29572 (2013).

31. Bradley ME, et al. Potent and efficacious inhibition of CXCR2 signaling by biparatopic nanobodies combining two distinct modes of action. Mol Pharmacol 87, 251–262 (2015).

32. Kim Y, Anderson JL, Lewin SR. Getting the “Kill” into “Shock and Kill”: Strategies to Eliminate Latent HIV. Cell Host Microbe 23, 14–26 (2018).

33. Crawford LB, et al. Human Cytomegalovirus US28 Ligand Binding Activity Is Required for Latency in CD34(+) Hematopoietic Progenitor Cells and Humanized NSG Mice. MBio 10, (2019).

34. Herbein G, Wendling D. Histone deacetylases in viral infections. Clin Epigenetics 1, 13–24 (2010).

35. Shin HJ, DeCotiis J, Giron M, Palmeri D, Lukac DM. Histone deacetylase classes I and II regulate Kaposi’s sarcoma-associated herpesvirus reactivation. J Virol 88, 1281–1292 (2014).

36. Shirakawa K, Chavez L, Hakre S, Calvanese V, Verdin E. Reactivation of latent HIV by histone deacetylase inhibitors. Trends Microbiol 21, 277–285 (2013).

37. Subramanian S, Bates SE, Wright JJ, Espinoza-Delgado I, Piekarz RL. Clinical Toxicities of Histone Deacetylase Inhibitors. Pharmaceuticals (Basel) 3, 2751–2767 (2010).

38. Duggan S. Caplacizumab: First Global Approval. Drugs 78, 1639–1642 (2018).

39. Poullin P, Bornet C, Veyradier A, Coppo P. Caplacizumab to treat immune-mediated thrombotic thrombocytopenic purpura. Drugs Today (Barc) 55, 367–376 (2019).

40. Bobkov V, et al. Nanobody-Fc constructs targeting chemokine receptor CXCR4 potently inhibit signaling and CXCR4-mediated HIV-entry and induce antibody effector functions. Biochem Pharmacol, (2018).

41. Qasemi M, Behdani M, Shokrgozar MA, Molla-Kazemiha V, Mohseni-Kuchesfahani H, Habibi-Anbouhi M. Construction and expression of an anti-VEGFR2 Nanobody-Fc fusionbody in NS0 host cell. Protein Expr Purif 123, 19–25 (2016).

42. Oliveira S, Heukers R, Sornkom J, Kok RJ, van Bergen En Henegouwen PM. Targeting tumors with nanobodies for cancer imaging and therapy. J Control Release 172, 607–617 (2013).

43. Behdani M, et al. Development of VEGFR2-specific Nanobody Pseudomonas exotoxin A conjugated to provide efficient inhibition of tumor cell growth. N Biotechnol 30, 205–209 (2013).

44. D’Huyvetter M, et al. (131)I-labeled Anti-HER2 Camelid sdAb as a Theranostic Tool in Cancer Treatment. Clin Cancer Res 23, 6616–6628 (2017).

45. Poole E, Reeves M, Sinclair JH. The use of primary human cells (fibroblasts, monocytes, and others) to assess human cytomegalovirus function. Methods Mol Biol 1119, 81–98 (2014).

46. Poole E, Walther A, Raven K, Benedict CA, Mason GM, Sinclair J. The myeloid transcription factor GATA-2 regulates the viral UL144 gene during human cytomegalovirus latency in an isolate-specific manner. J Virol 87, 4261–4271 (2013).

47. Weekes MP, et al. Latency-associated degradation of the MRP1 drug transporter during latent human cytomegalovirus infection. Science 340, 199–202 (2013).

48. Maussang D, et al. Human cytomegalovirus-encoded chemokine receptor US28 promotes tumorigenesis. Proc Natl Acad Sci U S A 103, 13068–13073 (2006).

49. de Wit RH, et al. CXCR4-Specific Nanobodies as Potential Therapeutics for WHIM syndrome. J Pharmacol Exp Ther 363, 35–44 (2017).

50. Heukers R, et al. The constitutive activity of the virally encoded chemokine receptor US28 accelerates glioblastoma growth. Oncogene 37, 4110–4121 (2018).

51. Slinger E, et al. HCMV-encoded chemokine receptor US28 mediates proliferative signaling through the IL-6-STAT3 axis. Sci Signal 3, ra58 (2010).

