## Supplemental Figures_De Groof, Elder et al_2020 for "Targeting the latent human cytomegalovirus reservoir with virus specific nanobodies"

**Supplementary Materials**

**Supplementary Figure 1**

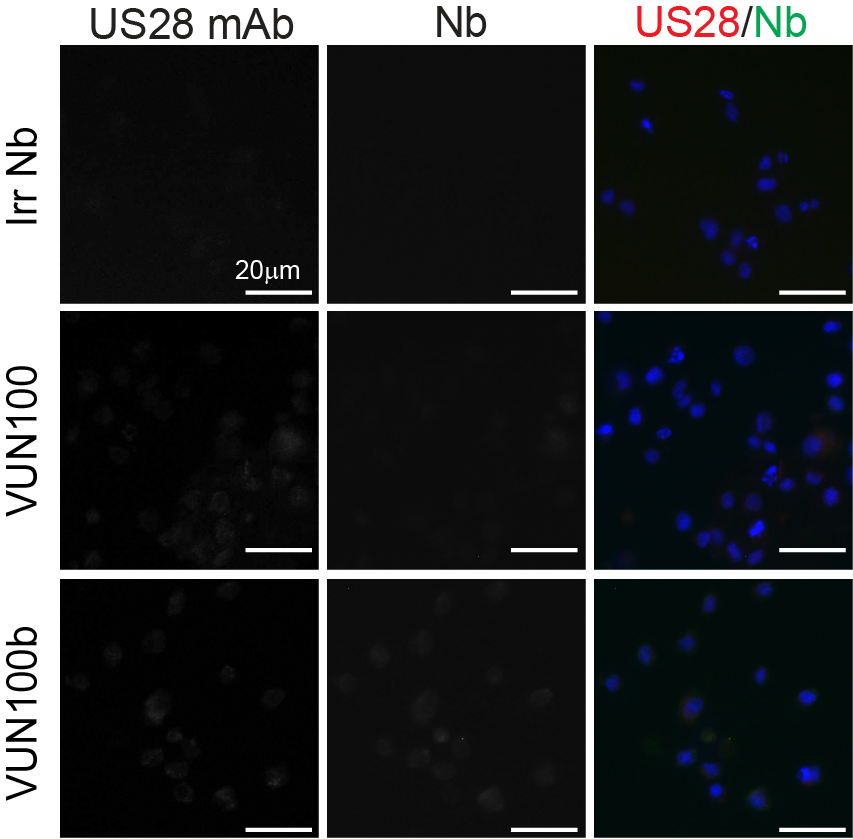

**Fig. 1. Binding of nanobodies to Mock transduced THP-1 cells.** Immunofluorescence microscopy of nanobody binding to mock transduced THP-1 cells. US28 was detected using an anti-US28 antibody (US28 mAb). Nanobody binding was detected using the Myc-tag and an anti-Myc antibody (Nb).

**Supplementary Figure 2**

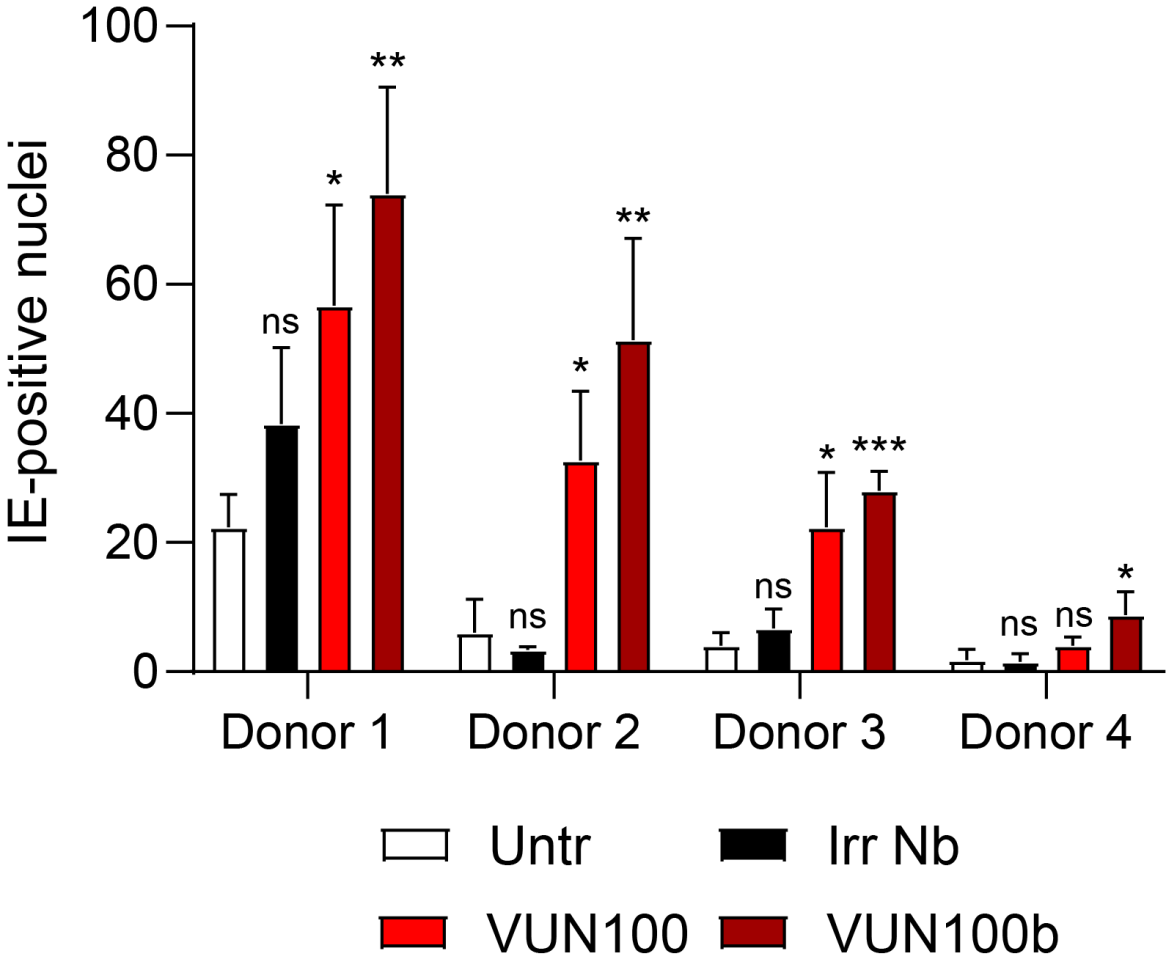

| **Donor** | **Untreated**  **(mean + S.D.)** | **Irr Nb**  **(mean + S.D.)** | **VUN100**  **(mean + S.D.)** | **VUN100b**  **(mean + S.D.)** |
| --- | --- | --- | --- | --- |
| **1** | 22 ± 4 | 38 ± 10 | 57 ± 13 | 74 ± 13 |
| **2** | 6 ± 4 | 3 ± 0 | 33 ± 9 | 51 ± 13 |
| **3** | 4 ± 2 | 7 ± 2 | 22 ± 7 | 28 ± 2 |
| **4** | 2 ± 2 | 2 ± 1 | 4 ± 1 | 10 ± 2 |

**Fig. 2. overview of immediate-early positive cells after nanobody treatment of latently infected CD14^+^ monocytes of 4 different donors.** CD14^+^ monocytes were isolated, infected with HCMV IE2-YFP and treated with an irrelevant nanobody (Irr Nb), VUN100 or VUN100b. IE2-positive nuclei were counted 6 days post infection. Data of four different donors is plotted as mean ± S.D. Statistical analyses were performed using unpaired two-tailed t-test and values were tested against untreated condition. ns, p > 0.05; *, p < 0.05; **, p < 0.01; ***, p < 0.001.

**Supplementary Figure 3**

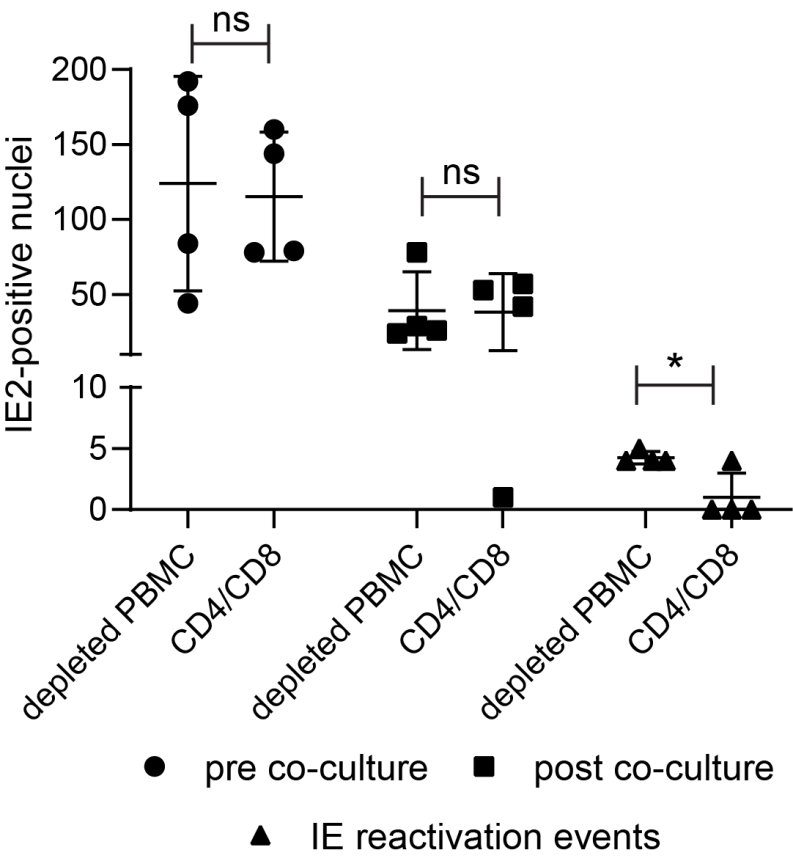

**Fig. 3. T cell/PBMC co-culture experiment of HCMV-infected PMA-pretreated CD14^+^ monocytes.** Counting of IE2-positive CD14^+^ monocytes before co-culture with CD4/CD8^+^ T cells or T-cell depleted PBMCs (pre co-culture), after 2 days of co-culturing of CD14^+^ monocytes and T cell depleted PBMCs or T cells (post co-culture) and after differentiation of CD14^+^ monocytes to mature dendritic cells (IE reactivation events). Cells were pre-treated with 20 ng/ml PMA before infection. Data is plotted as mean ± S.D.. Statistical analyses were performed using unpaired two-tailed t-test. ns, p > 0.05; *, p < 0.05.

**Supplementary Figure 4**

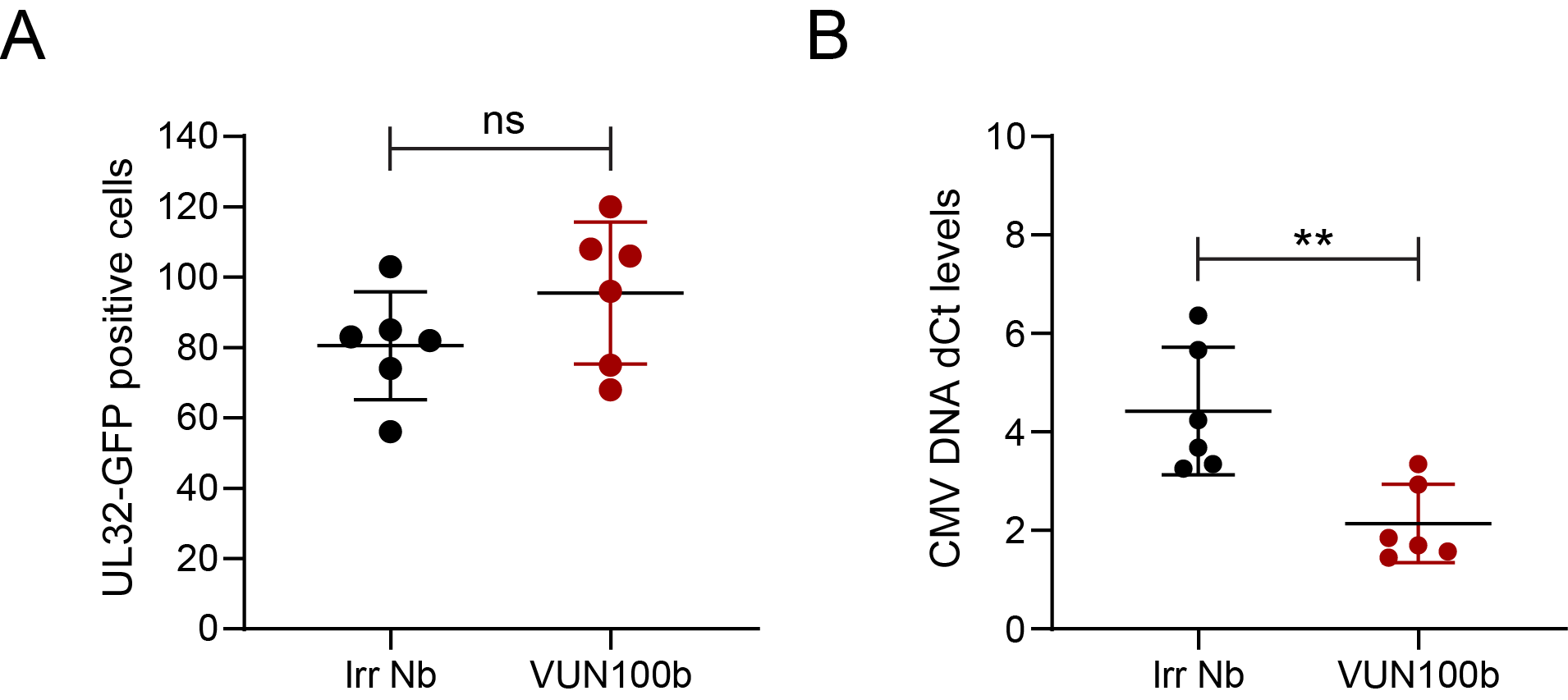

**Fig. 4. T cell co-culture experiment of CD14^+^ monocytes infected with HCMV UL32-GFP virus. A)** Counting of UL32-GFP positive CD14^+^ monocytes before co-culture with CD4/CD8+ T cells. CD14^+^ monocytes were treated with an irrelevant nanobody (Irr Nb) or VUN100b for 6 days post infection. **B)** Quantification of HCMV genomes after co-culture of CD4/CD8^+^ T cells, differentiation of CD14^+^ monocytes to mature dendritic cells and co-culturing of mature dendritic cells with Hff1 cells. Data is plotted as mean ± S.D.. Statistical analyses were performed using unpaired two-tailed t-test. ns, p > 0.05; **, p < 0.01.
